# Clearing the whey: Efficient inhibition and removal of chymosin from whey by de novo designed protein-based inhibitors

**DOI:** 10.64898/2025.12.23.695705

**Authors:** Darian Stephan Wolff, Casper de Lichtenberg, Josefine Gjelstrup Larsen, Carsten Andersen, Timothy Patrick Jenkins

## Abstract

Chymosins are indispensable as coagulants in cheese production. However, the residual activity of these enzymes in the whey by-product limits downstream processing for further food applications, e.g. infant milk formulations or protein powder for athletic purposes. To date, inactivation of coagulants and other enzymes is primarily achieved through heat inactivation (pasteurisation), which remains a challenging and energy-intensive task at an industrial scale and may have adverse effects on the quality. Here, we report the *de novo* design of small protein minibinders (miBds) that inhibit coagulant activity which marks one of the first industrial applications of generative binder design. Using RFdiffusion^1^, ProteinMPNN^2^, and AlphaFold2^3^, we generated and filtered over 2,000 backbone designs to a final panel of 63 candidates targeting bovine and camel chymosins. The first round of selected binders exhibited low-nanomolar K_i_^app 4,5^ values against chymosins originating from bovine, camel, and hedgehog, with reversible, pH-dependent dissociation profiles. Notably, we identified miBds specific to bovine and camel chymosin variants, as well as broadly cross-reactive including coagulants of fungal origin. Moreover, when immobilised, the miBds efficiently removed residual chymosin from whey. It was possible to recycle the miBd-coated resin repeatedly through treatment of the resin with high pH (> 11) or low pH (< 3) which emulates industry standard clean-in-place procedures. This dual functionality, of inhibition and reversible capture of chymosin, enables a versatile application for sustainable food production and precise control of proteases and can be extended to other enzyme classes in dairy.

## Main

Transformation of milk into a gel is arguably the most important step in cheese-making (Fig 1a). Here, the enzymatic coagulation of milk by chymosin (EC 3.4.23.4) is the chosen method for most modern cheeses^6^. Whey, the liquid by-product of cheese curd formation, is a versatile ingredient in the food industry. It contains high-quality proteins, lactose, vitamins, and minerals, making it valuable for applications ranging from infant formula and sports nutrition to functional beverages and baked goods^7,8^. Leveraging whey in food applications not only enhances nutritional profiles but also promotes sustainable use of dairy resources^7^.

**Figure 1.**
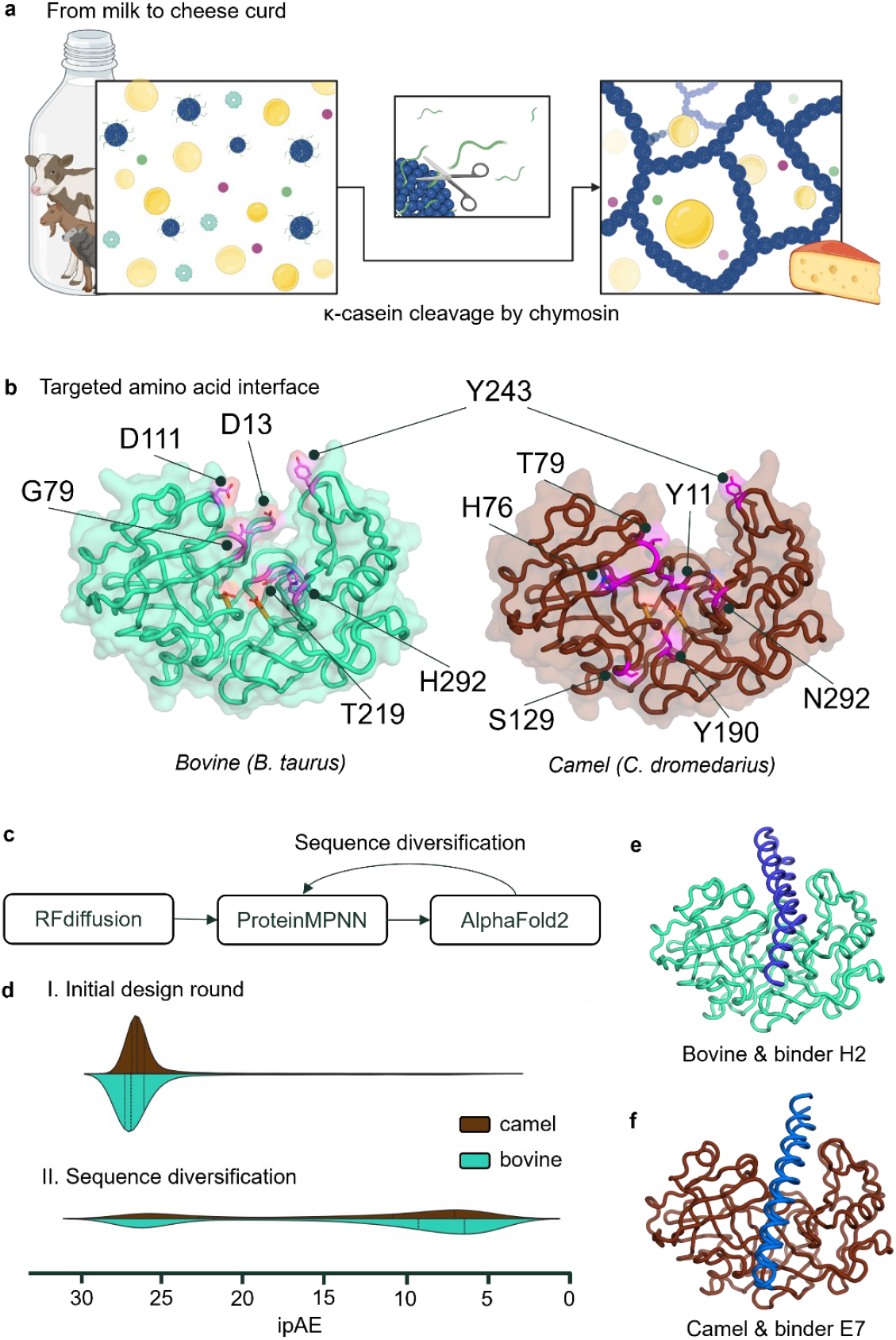
a, From milk to cheese curd – activity of chymosin. Milk contains casein proteins organised in micelles. When κ-caseins on the surface of the micelles are cleaved by chymosin (at the peptide bond between F105 and M106), the micelles lose their repulsion, coagulate and form a gel – the cheese curd. b, Targeted amino acid interface in bovine (left panel) and camel (right panel) chymosin. Surface exposed amino acids with outward facing side chains on each of the surrounding loop regions as potential contact points (defined as ‘hotspots’ in RFdiffusion) are indicated with their residue number in the polypeptides with PDB access codes: 4AA8 and 4AA9, respectively. c, Computational tool workflow used in the denovo design of the binders. d, Population distribution of the interface predicted alignment error (ipAE) of designs split into initial design round (I.) and after sequence diversification (II.) and campaign (camel in brown and bovine in petrol). e,f, Structure of the H2 bovine chymosin binder candidate (dark blue) binding to bovine chymosin (cyan) and E7 camel chymosin binder candidate (dark blue) binding to camel chymosin (brown), respectively. Structural visualisations in b, e, f, were obtained with pyMOL (version 3.1.6.1) and the backbone is displayed in ‘cartoon tube’.

Despite the advantages whey holds, the residual chymosin-driven coagulant activity in the whey fraction poses a substantial hurdle for high-value downstream applications. As such, industrial processes often employ pasteurisation (e.g., 15 s at 72 °C) to remove residual enzyme activity. This is an energy-intensive step that denatures valuable whey proteins and does not fully address enzyme carry-over across all processing schemes. Furthermore, during the holding time between cheese curd formation and pasteurisation the enzyme remains active and may produce off-flavours.

Consequently, there has been an effort to explore alternative modes of chymosin deactivation such as inhibition by ligand binding. Pepstatin, first identified in the culture filtrate of various actinomyces by Umezawa et al. (1970)^9^, is a well-studied inhibitor of aspartyl proteases such as chymosin. The statine moiety, a non-proteinogenic amino acid, plays a pivotal role in competitive inhibition by occupying the catalytic aspartic acid residues of the enzyme.^10^ This disrupts the coordination of the water molecule required for nucleophilic attack and prevents hydrolytic cleavage of the substrate, κ-casein^10^. Yet, its industrial potential for coagulant removal is limited. A key issue is pepstatin’s lack of specificity, since it strongly inhibits porcine and human pepsin^11–13^ and is effective against most acidic proteases^11–13^, raising pharmacological and regulatory concerns especially for the high-quality and safety standards required in infant formula. Moreover, its low solubility in aqueous solvents^13^ render it unsuitable for industrial food applications.

Recent breakthroughs in de novo protein design offer promising avenues to address this issue. Protein design tools including RFdiffusion^1^ and ProteinMPNN^2^, as well as protein structure prediction tools such as AlphaFold^3^ (AF) have enabled the creation of custom protein minibinders (miBds) with nanomolar affinity to targets with high specificity or controlled cross-reactivity. While these binders so far primarily have been used for therapeutic neutralisation of pathogenic proteins^15–17^, their potential to inhibit and/or capture proteases for downstream removal remains largely unexplored in industrial biotechnology. We reasoned that selective occlusion of the active site in chymosin, would lead to inhibition of its function. As a benefit of the specific inhibition of only targeted coagulants, foremost bovine chymosin, the inhibitor and the targeted chymosin can remain as an inactive complex in the product without adverse effects to the whey consumer. In another aspect, the strong binding affinity of these miBds offer an appealing method for enzyme removal from whey filtering through a matrix with immobilised binders. In this case, broad cross-reactivity to a defined array of coagulants is beneficial as it reduces the number of different miBds required to retain all the coagulants in the portfolio of modern cheese manufacturers. Both the in-solution inhibition and the removal of chymosin eliminate the need for a high-temperature processing step and enhance product quality.

### Design of chymosin minibinders

We first sought to create compact de novo proteins capable of sterically blocking the active-site cleft of either bovine or camel chymosin without perturbing the overall fold. Crystal structures of the bovine (*Bos taurus*) and camel (*Camelus dromedarius*) chymosin (PDB access codes: 4AA8 & 4AA9, respectively) reveal three surface-exposed loops (residues 76–85, 235–245 and 283–291) framing the catalytic dyad (Asp 34 and Asp 216)^18^. Using RFdiffusion, we generated 1,500 backbone-only scaffolds per target, every trajectory conditioned to engage one of those loops (Fig 1b). Each scaffold was passed through ProteinMPNN to propose two sequences; binder-target-complex models were predicted with AF2 (Fig 1c). Subsequently, designs were retained if they showed an interface predicted alignment error (ipAE) below 15 Å, a predicted local distance difference test score (pLDDT) of above 90, and no steric clashes; this yielded 238 candidates for bovine and 86 for camel chymosin (Fig 1c and d). The lower hit rate on the camel enzyme reflects the flexibility of the N-terminal region^18^, which is integrated in the central antiparallel better sheet in bovine, but not in camel chymosin.

To balance the two target panels, we carried out a second round of RFdiffusion (1,500 additional backbones) with four sequences sampled per backbone for camel chymosin. After AF2 filtering, 100 new candidates passed the same cut-offs. We then applied a final sequence-diversification step to all selected bovine and camel chymosin binder designs, resampling each candidate with ProteinMPNN to generate 32 variants and imposed stricter criteria of an ipAE below 6 Å and pLDDT above 92 (Fig 1d). From this, we distilled a final set of 42 miBds against bovine chymosin and 21 against the camel enzyme. All selected designs are 50–100 amino acids long and adopt a two-helix bundle that cradles the frontal loops and plugs the catalytic groove of the chymosin (Fig 1e and f).

### Identification of chymosin inhibitors

To assess if the designed binders (miBds) inhibit chymosin, we first expressed the final design set of 63 candidates in *Bacillus subtilis* and purified them via Ni-NTA affinity chromatography. Genome integration was successful for 58 miBds. Despite variable expression (Fig. S1), all 58 purified samples were screened for inhibition of bovine chymosin in a milk-clotting assay. Fifteen miBds reduced bovine chymosin activity, with miBd H2 showing the strongest effect (>95% reduction; Fig. 2a). Notably, some potent inhibitors (e.g., B4) originated from samples with minimal detectable expression.

**FIGURE 2.**
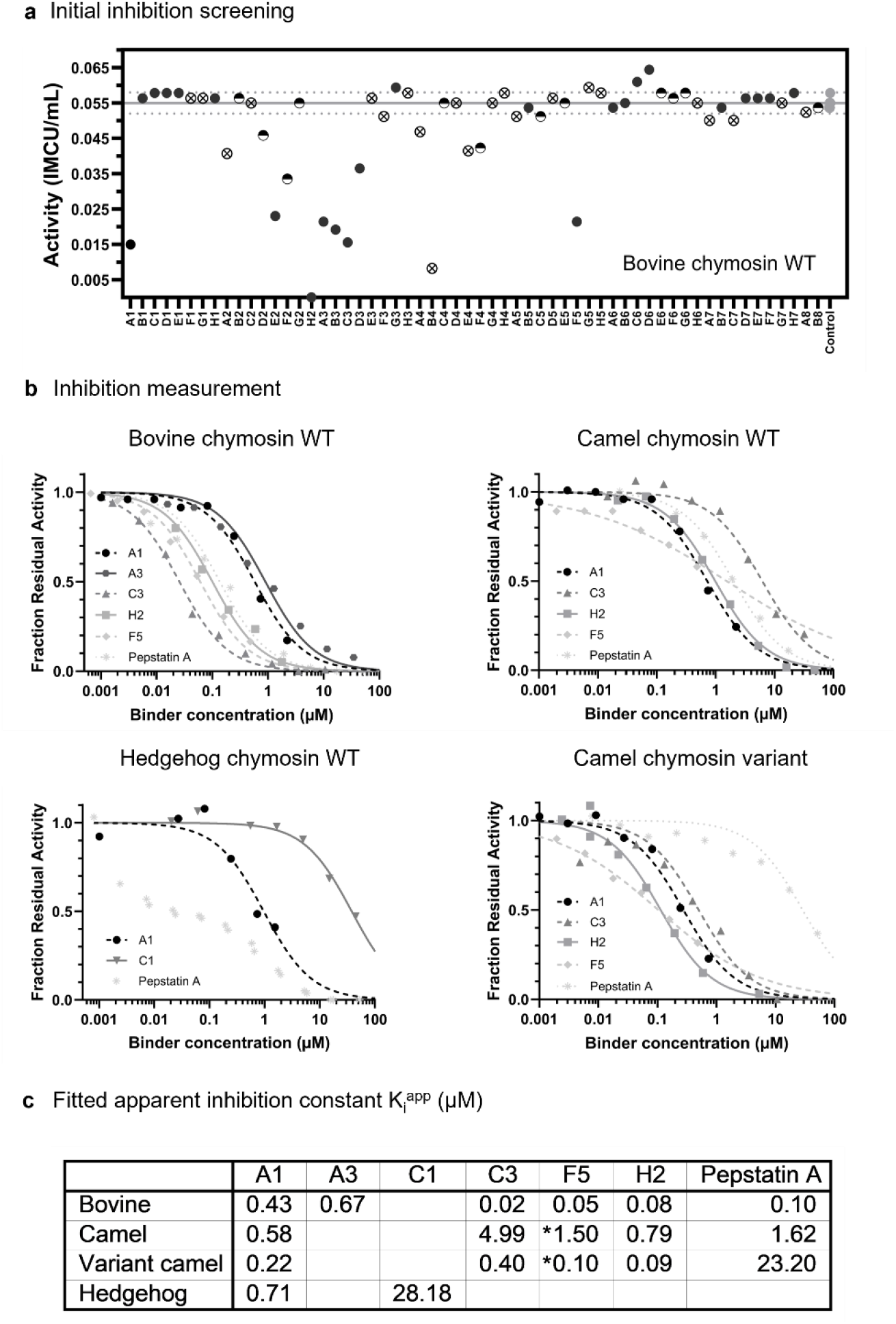
a, Initial inhibition screening of the (58) expressed and purified binder designs (x-axis) towards bovine chymosin. The miBds were added in the same volume and not adjusted for concentration. An indication of the concentration was obtained by their visibility on SDS-PAGE gel. Dark (full) circles symbolise miBds with clearly visible expression. Similarly, half-filled circles were barely visible and crossed circles were not visible on the gel (Full gel picture in SF. 1). b, Quantitative inhibition measurement. For the miBds A1, A3, C1, C3, F5, and H2 the inhibition capacity was quantified and fitted using the Morrison approximation for determination of K_i_^app^ under tight binding conditions (Eq. 1) through the GraphPad Prism 10.2.1 suite. To benchmark the miBds pepstatin A was tested under the same conditions. c, Fitted K_i_^app^ values (µM), *F5 values for camel coagulants were derived as IC50 in a four-parameter logistic curve. The graphs for a,b were prepared with GraphPad Prism 10.2.1 suite.

Next, while bovine chymosin presents the most commonly used coagulant in cheese production, we extended the panel to include further commercially relevant or putatively relevant chymosin variants originating from camel and hedgehog chymosin. MiBds H2 and F5 consistently inhibited all tested variants, while B4 preferentially targeted bovine and wild-type camel chymosin (>80% and >60% reduction, respectively) but was less effective against the camel variant (<40%). MiBd A3 displayed selective inhibition of bovine chymosin and the camel variant (Fig. S2).

Based on these results, we characterised the inhibition kinetics of the most promising miBds and selected the six top candidates (miBds A1, A3, B4, C3, F5, H2) as well as miBd C1 since it revealed highly specific inhibition of hedgehog chymosin. Apparent inhibition constants (K_i_^app^) were determined using established kinetic models^4,5^ and ranged from 28 µM to 20 nM, which confirmed diverse and strong inhibitory profiles (Fig. 2b,c,d). MiBd C3 exhibited the highest potency of any miBd against bovine chymosin (K_i_^app^ = 20 nM). For camel chymosin, several miBds (A1, C3, F5, H2) showed markedly improved inhibition of the camel variant compared to wild-type (up to 15-fold difference), indicating variant-specific binding preferences.

To better unravel how the miBds performed compared to the current state-of-the-art chymosin inhibitor, pepstatin was tested under identical conditions. While pepstatin inhibited bovine chymosin (K_i_^app^ 100 nM ) comparable to some miBds (Fig. 2c), it was significantly less effective against the camel wild type and variant (K_i_^app^ = 1.62 µM and 23 µM, respectively), highlighting the superior performance of miBds such as A1, C3, F5, and H2 (K_i_^app^ for camel variant 90-400 nM).

Finally, since both the specificity to bovine chymosin and the cross-reactivity to a broad spectrum of commercially available coagulants are vital for either in-solution inhibition or broad-spectrum retention, we extended the panel of coagulants to include bovine variants of other vendors, bovine pepsin, and coagulants of fungal origin, that also represent a large share of the coagulant market (Fig. 3a). To include almost all miBds that revealed inhibition capacity to at least one of the chymosins in the initial activity, we broadened the panel of miBd and included the miBds B1, B2, B3, C2, E7 at the highest available concentration. While each of the miBds A1, B4, C3, F5, and H2 inhibited all bovine variants, especially miBds A1 and F5 revealed little (>80% residual activity) to no inhibition effect on bovine pepsin. MiBds A3 and B3 reduced the activity of wild type bovine, but not or only partially of the bovine variants and bovine pepsin. In consequence, miBds A1, A3, B3, and F5 might be considered suitable for in-solution inhibition. Beyond the specific recognition of bovine chymosin but not bovine pepsin, miBd A1 revealed strong inhibition of the fungal coagulant mucorpepsin. Other miBds revealed a broad spectrum of inhibition extending to fungal coagulants. MiBds B2, B4, and H2 broadly inhibited bovine and camel chymosin wild types and variants as well as coagulants of fungal origin, potentially making them better suited for the immobilisation of a wide array of coagulants on a filter device. Overall, these findings underscore the broad inhibitory spectrum and specificity of the miBd library, exceeding initial design expectations.

**FIGURE 3.**
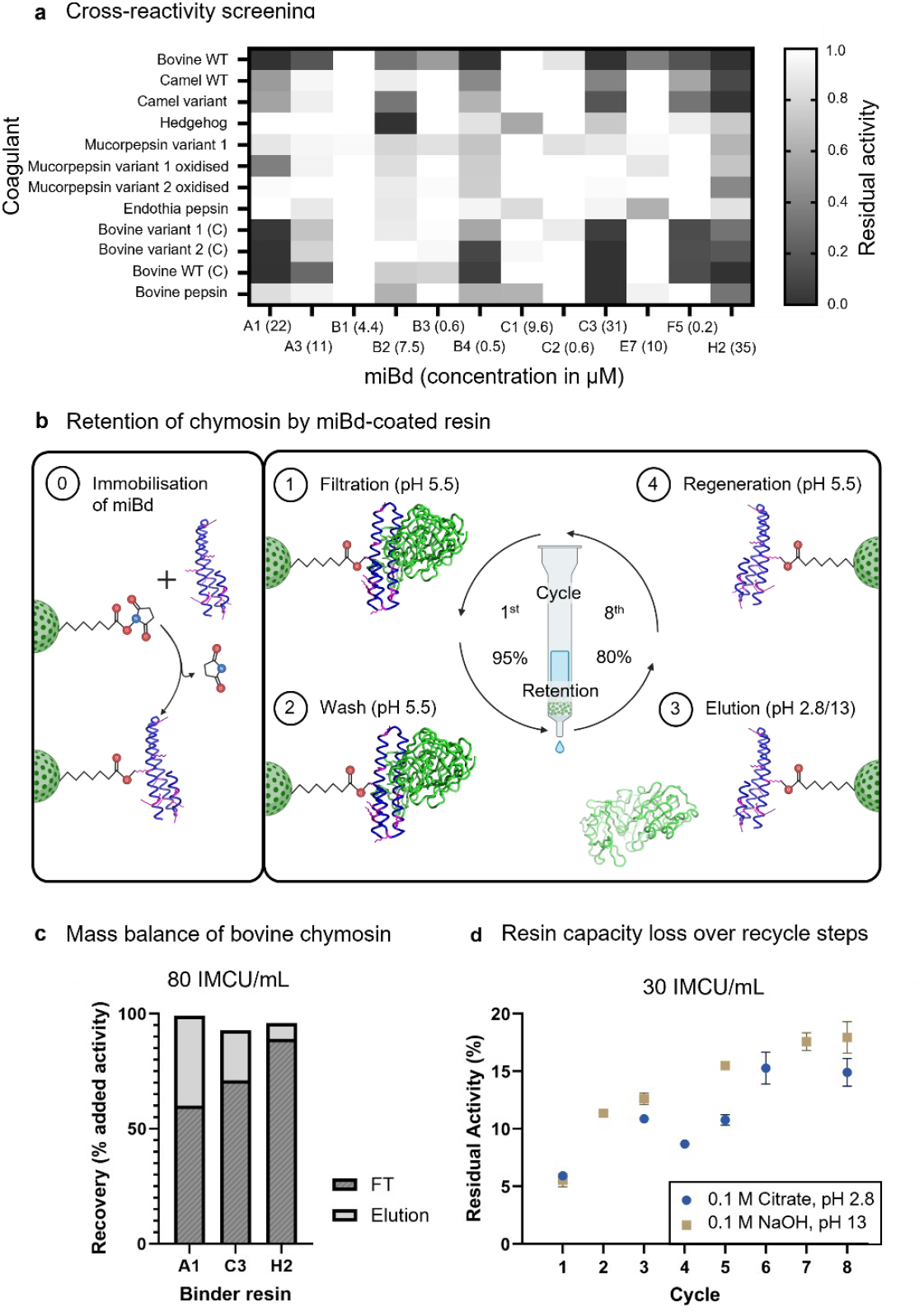
a, Heat map displaying the residual activity of a broad panel of coagulants (y-axis) in presence of miBds (X-axis). The panel of coagulants included in the assays were bovine and camel chymosin variants, hedgehog chymosin, coagulants of fungal origin (Mucorpepsin and their oxidised variants, Endotheia pepsin), bovine variants obtained from vendors other than Novonesis A/S (Bovine variant 1 is Maxiren EVO from DSM, Bovine variant 2 is Maxiren XDS from DSM and Bovine WT is chymostar from IFF), and bovine pepsin. b, Schematic display of the retention experiment. First, the relevant miBds (A1, C3, & H2) were covalently crosslinked to N-hydroxysuccinimidyl Sepharose matrix. Next, whey containing high concentrations of chymosin was applied to the gravitational flow columns. The retained chymosin was eluted at either high or low pH. After regeneration at application pH, the matrix could be subjected to a new filtration cycle. As in Fig. 1b & e, the bovine chymosin is depicted in green, while the miBd shown in dark blue. The side chains of lysine in the miBd can serve as anchor residues for covalent crosslinks to the matrix (highlighted in pink). The images for panel b were generated and arranged using Biorender.com. c, Mass balance of chymosin based on residual activity before (control) and after filtration. The chymosin after filtration is divided into the chymosin that remained in the whey fraction (FT, flow through), depicted in in dark grey, and the fraction of chymosin that was previously retained and eluted at low pH (light grey). d, The matrix was re-used several times (8 cycles in total) and the residual chymosin activity in the filtered whey fraction measured for each cycle. The graphs in panel a, c, and d, were plotted using GraphPad Prism 10.2.1 suite.

### Retention and release of chymosin in whey by miBd-coated resin

We evaluated if the miBds covalently linked to a solid resin could serve as a filter medium to remove chymosin from whey. Early, near-complete removal of chymosin from the whey fraction in the dairies by simple filtration is ideal for fast preservation of the nutritional and functional value of the whey and alleviates the need for pasteurisation at a later time at whey processing plants. Furthermore, it is essential that the filter matrix can be efficiently recycled by elution of the bound chymosin (ideally without removing it from the process stream). Therefore, three of the most relevant miBds, based on expression yields, potency and broad specificity (A1, C3, & H2) were covalently crosslinked to N-hydroxysuccinimidyl Sepharose (Fig. 3b). We then determined the capacity of each resin for removing chymosin under gravitational flow, by comparing the residual activity of chymosin in the filtrate and eluate (with 0.1 M citrate pH 2.8) to the chymosin before filtration (Fig. 3c). For all three miBds, more than 90% of the original chymosin activity was recovered when summing the activity measured in the filtrate and the eluate, and recovery was nearly complete for miBd A1. This demonstrates that chymosin bound to the resin could be efficiently released during elution. Among the tested binders, miBd A1 showed the highest ability to capture bovine chymosin. When whey containing 80 IMCU/mL chymosin (approximately 9.1 nM) was passed through the A1-coated resin, about 40% of the initial chymosin activity was recovered in the eluate. Based on these measurements, 10.5 nmol of immobilised miBd A1 retained 3.6 nmol of bovine chymosin. This corresponds to an efficient stoichiometry, requiring only a 2.9-fold molar excess of immobilised binder relative to chymosin in the whey fraction.

After establishing the general ability to efficiently elute the bound chymosin, we evaluated the reusability of miBd-coated resin under harsh and standard industrial cleaning/elution conditions with low (as before, 0.1 M citrate, pH 2.8) and high pH (0.1 M NaOH, pH 13). At a concentration of 30 IMCU/mL of bovine chymosin in the whey fraction, the same quantity of A1-coated resin retained 95% of bovine chymosin in the initial round. Independent of the elution condition, high or low pH, the resin capacity decreased to 90% retention within a single recycling step. Within the following six filtration and elution cycles the resin capacity decreased at substantially slower rate. Between the 2^nd^ and the 8^th^ cycle, the miBd-coated resin lost less than 10% retention capacity. Consequently, after the 8^th^ cycle >80% of the chymosin in the applied whey was retained. In result, the miBd-coated resin was found to be reuseable and compatible with harsh conditions, which likely decreases replacement costs and increases the value-proposition in industrial applications of whey filtration. Since there was no attempt to optimise e.g., the orientation of the miBd A1 nor to improve the binding affinity of the miBd itself, prospective optimisation will likely lead to higher retention capacity and/or longevity of the miBd-coated resin.

### Characterisation of binding affinity and interaction

Since the miBds’ binding affinity regardless of the capability of inhibition is the main reason for the retention capacity of the miBd-coated resin, we investigated the binding properties of miBds A1, C3, and H2 (Fig. 4). Similar to the filtration experiment, we immobilised each of the miBds on antibody-coated biosensors and measured the binding affinity towards chymosin in solution. Using biolayer interferometry (BLI) we obtained affinity measurements beyond background signal for 7 out of 9 combinations of miBds A1, H2, and C3 towards bovine, camel, and camel variant chymosin (Fig. S3 c-f). Only for the combinations of miBd C3 towards the camel chymosins the binding affinity could not be decisively resolved. Remarkably, when comparing the dissociation of chymosins from the miBd-coated biosensor, a striking difference in the dissociation rates between miBd A1-and miBd H2-coated biosensors was observed. Regardless of the origin or variant of chymosin the dissociation rates were between 1.5-and 10-fold lower for miBd A1 (_kdis_ ranged from 9.9×10^-4^ to 4.1 ×10 ^-3^ s^-1^), and miBd C3 (towards bovine chymosin: k_dis_ 8.6x10^-4^ to 4.1x10^-3^ s^-1^) in comparison to miBd H2 (k_dis_ 6.1x10^-3^ to 1.1x10^-2^ s^-1^). In consequence, the lower dissociation rate likely allowed for the comparatively higher retention capacity of miBds A1 and C3 in the filtration of chymosin from whey, when compared to H2. These differences in dissociation rates led to a twofold worse, but nevertheless nanomolar binding affinity of miBd H2 in comparison to miBd A1 towards both wt bovine and camel variant chymosin (K_d_ values towards bovine chymosin: 38, 63, and 83 nM for miBd A1, C3 and H2, respectively) (Fig. S3c–f and h).

**FIGURE 4.**
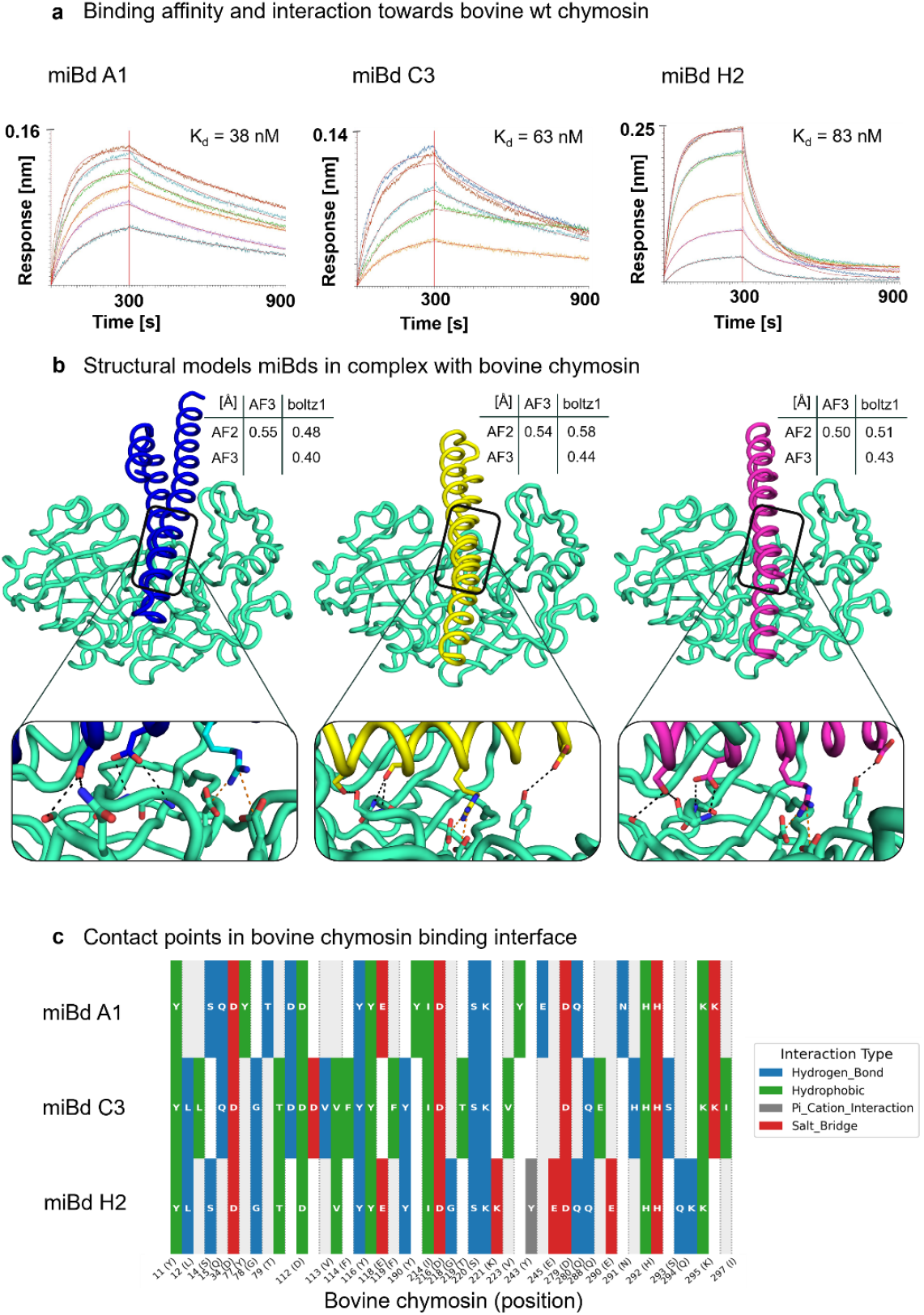
a, Measuring binding affinity. Association (first 300 seconds (s)) and dissociation (600 s) curves of various concentrations of bovine wt chymosin in solution towards immobilised miBd (A1, C3, H2) on biosensors. The red solid lines, laid-over raw data, represent a 1:1 ligand model fit and were used to derive dissociation constant for each combination of miBd and chymosin. b, Structural models of bovine wt chymosin (turquoise) with each of the miBd A1 (blue), C3 (yellow), and H2 (purple) derived from AF3. RMSD of all atoms except hydrogens between structural models from boltz-1, the initial AF2, and AF3 structures are provided in an adjacent table. The interactions between the miBd and the chymosin were obtained by using a customised version of PLIP^21^. A selection, non-exhaustive number of hydrogen bonds are displayed in dashed black lines. The salt bridge between chymosin residues D34 and D279 and the arginine of the miBd is depicted in an orange dashed line. c, Contact points on the chymosin surface (y-axis) bound by each of the miBd A1, C3, and H2. The light grey and white tiles are identical, symbolise the white space, and are different in shade to distinguish adjacent target positions. For the visualisation of panel c the python package matplotlib was used. The structural visualisation were obtained with pyMOL (version 3.1.6.1) and the backbone is displayed in ‘cartoon tube’.

To structurally explore the binding interface, we re-predicted the co-complexes of each miBd with bovine and camel chymosin using sequence input alone. The structures obtained from boltz-1 and AF3 were nearly identical to each other and to the initial structural models derived from AF2. For any combination of miBd with bovine or camel wild-type chymosin, the root mean square deviation (RMSD) of all atoms, excluding hydrogens, for the aligned structural predictions from AF2, AF3, and boltz-1 did not exceed 1.5 Å. This level of agreement between prediction models, and the precise match of modelled and experimental solved structures of de novo binder-antigen complexes in previous cases^16,17^, gave us sufficient confidence to proceed with all further analyses using AF3-derived structure predictions. Overall, we found that the miBds were predicted to be directly embedded in the active site (Fig 4a), surrounded by three surface-exposed loop regions (residues 76–85, 235–245, and 283–291) where the majority of miBds, regardless of the chymosin type, bound (Data S3 and S4). Additionally, most miBds interacted with residues 112-116 and 190, previously defined as putative contact points (‘hotspots’) (Fig 1b) in the bovine design campaign. Notably, many miBds are predicted to form salt bridges with the catalytically active aspartic residues (D34 and D279).

In terms of inhibition, (1) the potential interaction with D34 and/or D279 could disrupt the coordination of the activated water molecule, which is crucial for chymosin’s catalytic activity^19^. Furthermore, (2) the strong binding within the active site pocket prevents any K-casein annealing. Pepstatin establishes similar interactions with the aspartic residues. A previously resolved crystal structure (PDB access code: 4AUC^20^) of the pepstatin-bovine chymosin complex suggests one or more hydrogen bonds between the hydroxyl group of statine (pepstatin, position 4) and D34 and D279. In our structural predictions, miBds A1, C3, and H2 share several interactions with chymosin (Fig 4a); for instance, all three miBds are predicted to bind to the aspartic residues with an arginine (Fig 4b,c).

Together, kinetic and structural analyses indicate that miBds inhibit chymosin by active-site occlusion and stabilise capture on resin through slow dissociation. The consistent engagement of the designed loop epitope and predicted contacts to the catalytic aspartates provide a mechanistic basis for inhibition, whereas differences in retention under flow were largely explained by the dissociation rate rather than by equilibrium affinity. These results highlight kinetic optimisation as a key axis for translating these de novo binders from in vitro inhibition to robust performance in complex process streams.

## Discussion

De novo protein design has transformed the development of compact binders for therapeutic targets^15– 17,22–24^, but its use in industrial biotechnology has been limited by a lack of demonstrably functional designs that operate reliably in complex process streams. Here we show that generative design can yield miBds that not only potently inhibit the residual activity of chymosin in whey but also capture and remove the enzyme under conditions fully compatible with dairy processing. Residual chymosin poses a central barrier to the use of whey in high value applications such as infant formula, specialised nutrition, and high purity protein ingredients, because even low levels of protease activity degrade whey proteins and alter flavour and functionality. A selective, reusable and food-grade method for controlling coagulant activity therefore addresses a longstanding unmet need in dairy downstream processing.

Despite a modest size of the design library, the miBds generated here showed a comparatively high functional hit rate, with 18 of 63 designs inhibiting at least one coagulant. Several reached low nanomolar apparent inhibition constants against bovine chymosin, and multiple designs retained substantial activity against camel and fungal chymosins. Importantly, some miBds displayed minimal inhibition of pepsin. This discrimination is critical because broad inhibitors such as pepstatin, although potent, act across acidic proteases and carry pharmacological liabilities that limit their use in food-grade applications, especially those intended for infant consumption. The ability to generate protein-based inhibitors that match or exceed the potency of pepstatin while avoiding its off-target activity represents a meaningful advance for industrial enzyme control. The striking asymmetry in successful designs between the bovine and camel campaigns reflects underlying differences in designability. Bovine chymosin presents a rigid, well defined active site environment suitable for generative modelling, whereas camel chymosin features increased loop mobility^18,25,26^, substitutions within the substrate binding region^27^ and glycosylation at Asn291^27,28^. These structural features, not fully captured by the available crystal structure^18^, likely reduce model fidelity and explain the lower number of successful camel-focused designs. This suggests that future workflows incorporating flexible loop modelling or explicit glycan handling may expand the range of enzymes that can be targeted with confidence.

Beyond soluble inhibition, an essential industrial requirement is the ability to remove residual chymosin from whey. Immobilised miBds A1, C3 and H2 efficiently captured bovine chymosin, with nearly complete recovery upon elution, and maintained more than 80 percent of their capacity after repeated regeneration using both low and high pH conditions. These regeneration tolerances are central for integration into standard clean-in-place cycles and ensure operational feasibility without introducing new process constraints. The kinetic measurements provide a mechanistic rationale for the observed performance: binders with slower dissociation rates exhibited more effective retention under flow, even when equilibrium affinities were similar. This highlights off-rate optimisation as a key axis for improving binder performance in future design cycles.

Current reliance on heat inactivation through pasteurization constrains the development of novel coagulants. Candidates with desirable proteolytic activity or unique flavor profiles are often discarded or engineered to ensure heat sensitivity. By enabling inhibition or capture of coagulants, the developed binders eliminate this constraint, allowing cheesemakers to fully explore new coagulant options. Moreover, the de novo design framework offers rapid adaptability to emerging coagulants beyond the current binder set. Taken together, these results demonstrate that generative protein design can produce selective inhibitors that both suppress coagulant activity in whey and enable its removal in an energy-efficient, reusable manner. By avoiding the broad protease inhibition and regulatory challenges associated with small molecule inhibitors, and by circumventing the protein damage caused by heat inactivation, designed miBds offer a route to generate whey streams suitable for the most stringent applications, including infant formula. More broadly, this approach provides a general framework for deploying de novo proteins as functional processing aids and suggests that designed binders could be extended to other proteases or enzyme classes central to food and industrial biotechnology.

## Supporting information

Supplementary data (including sequences and structures of miBds)

## Acknowledgements

We are grateful for the invaluable assistance and diligence of Sadi Sivanendiran and Ulla Wahlers in the functional inhibition assays. Furthermore, we are indebted for the substantial assistance of Julien Menard in the draft of the related patent application. We appreciate the careful commentary of Isabela Rister Portinari Maranca, Christian Jäckel, and David Jenkins.

## Author contributions

(1) C.L., D.S.W., C.A., and T.P.J. conceived the project. (2) T.P.J. and C.A. supervised the work. (3) T.P.J., D.S.W., and C.L. wrote and reviewed the manuscript. (4) D.S.W. designed and filtered the miBd designs. (5) D.S.W. expressed and purified the miBds. (6) C.L. and J.G.L. performed all inhibition studies and (7) C.L. analysed and fitted the inhibition data. (8) C.L. and D.S.W. conducted the whey filtering. (9) D.S.W. made the binding affinity measurements. (10) All authors reviewed and approved the final manuscript.

## Data availability

All data needed to evaluate the conclusions in the study are available in the main text or the supplementary materials. Furthermore, we provided the structural models (from the AF2 filtering step and the sequences for miBds that were selected for the experimental assessment in the supplementary data (S1). Raw and fitted data of the affinity and the inhibition analyses can be found in the supplementary materials (S2). The code for the binder design was published elsewhere and available under BSD license (https://github.com/RosettaCommons/RFdiffusion)^1^. The visualization of contact points on the surface of chymosin was based on existing and public available code (PLIP 2025)^20^ under GPL-2.0 license.

## Methods

### De novo chymosin binder design using RFdiffusion

The crystal structures of the bovine (*B. taurus*) and camel (*C. dromedarius*) chymosin (PDB access codes: 4AA8 & 4AA9, respectively) were used as input to RFdiffusion. Based on the observation that the aspartic dyad, holding the activated water molecule, within the active site cleft were flanked by several loops, we deliberately defined surface exposed amino acids with outward facing side chains on each of the surrounding loop regions as potential contact point (Fig. 1b). Initially, we generated 1,500 diffused backbone designs per target using RFdiffusion with a length between 50-100 amino acids and employed ProteinMPNN to design four amino acid sequences per backbone. The resulting backbone libraries were assessed by AF2 with initial guess. Applying an interaction predicted alignment error (ipAE) of the chymosin:binder complex below 15 Å and the predicted local distance difference test (pLDDT) with a score above 90 resulted in a filtered subpopulation of 238 miBd designs for bovine chymosin and 86 designs for camel chymosin. To balance the number of selected designs originating from the bovine and camel chymosin, the design campaign of binders towards camel chymosin was repeated. Thereby, additional 1,500 backbones with four sequences each were generated, of which 100 new miBd designs for camel chymosin passed the same filter criteria. Following the initial selection, the subset of selected miBd designs was diversified in sequence. At the default temperature of t=0.1, 32 sequences were generated for each selected design using ProteinMPNN and reinterpreted into structures by AF2. Employing a more stringent cut-offs for ipAE <6 Å and pLDDT >92, a final set of 42 bovine and 21 camel chymosin miBd were subjected to experimental validation. All manual inspection for structures and figures of miBds for the manuscript have been made using pyMOL under academic license^29^.

### Transformation and expression of chymosin miBds

For the present study the miBds have been expressed in *Bacillus subtilis* A164Δ5 derivative (Widner et al. 2000^30^) and purified employing His-tag affinity purification. Briefly, the amino acid sequence of the miBd designs were complemented with an N-terminal secretion signal peptide from termamyl (‘MKQQKRLYARLLTLLFALIFLLPHSAAAA’) and a C-terminal 6xHis-tag (‘HHHHHH’). The de novo binders were ordered as synthetic gene fragments and cloned into the expression vector pDG268Δneo (Widner et al. 2000^30^). A recombinant clone containing the integrated expression construct was identified and grown in liquid culture medium (Widner et al. 2000^30^). For the verification of the correct integration of the miBd-encoding fragments the transformed Bacillus clones were grown on agar plates containing 6 mg/L chloramphenicol. For each miBd, four clones were selected and sequenced. For 58 out of 63 miBds the correct miBd-encoding DNA was identified. The miBds for which no correct sequence was obtained, were: (1) chymosin_bovine_miBd_NAN_6_51_2_5, (2)chymosin_bovine_miBd_NAN_9_84_1_22, (3) chymosin_bovine_miBd_NAN_12_43_2_2, (4)chymosin_bovine_miBd_NAN_13_4_0_23, (5) chymosin_camel_miBd_NAN_23_56_2_1. All information for these and the successfully cloned designs including the in silico scores for filtering, the structures, the amino acid and DNA sequences can be found in the supplementary (S1).

### Purification of chymosin miBds

For miBd purification, 250 mL of the sample containing cultures was subjected to centrifugation (10,000 x g for 15 min., Sorvall RC 6 plus, 46915 ThermoFisher Scientific), and the supernatant was applied to single-use columns for immobilised metal affinity chromatography (His GraviTrap columns, 11003399, Cytiva) following the supplier’s protocol. Subsequently, the samples were desalted employing disposable PD-10 columns packed with Sephadex G-25 resin (17085101, Cytiva) and eluted in 5 mL of buffer solution (50 mM MES, 50 mM Sodium acetate, 100 mM NaCl, pH 5.5). The concentrations of binders were determined utilising the Qubit Protein assay kit (Q33211, Invitrogen/ThermoFisher Scientific). Guided by the product manual, reagent buffer and protein assay buffer were mixed in the ratio 1:199, and 198 µL of the resulting reaction buffer were incubated with 2 µL of protein sample. For the standard curve 10 µL of each of the three supplied standards were incubated in 190 µL reaction buffer. After the recommended 15 min. incubation, a standard curve was established and sample concentration determined by measuring the standards and samples on the Qubit 4 fluorometer (Q33238, Invitrogen/ThermoFisher Scientific).

### Binding affinity measurements

The affinity measurements of miBd A1, B4, C3, F5, and H2 towards bovine, camel and camel variant chymosin were conducted on an Octet Red96 (ForteBio) instrument using biolayer interferometry (BLI).

#### A1 - Bovine Chymosin

The affinity measurements of miBd A1 towards bovine chymosin were conducted using HIS2 biosensors (Sartorius, 18-5114). All dilutions of antigen and miBd were prepared with MES acetate buffer without Tween 20 (MES-nT) [50 mM MES, 50 mM sodium acetate, 150 mM NaCl, pH 5.5]. Prior to the experiment the HIS2 biosensors were incubated for at least 10 min in MES-nT. For the binding assay, we used the miBd at 200 nM in MES-nT for immobilisation on HIS2 biosensors and measured the bovine chymosin as analyte in solution in seven concentrations ranging from 200 nM to 20 nM [200, 140, 100, 80, 60, 40, and 20 nM, respectively]. Experiments were performed at 30 °C, and data were fitted with a 1:1 binding model.

#### A1 - Camel Chymosin (wild type and variant)

The affinity measurements of miBd A1 towards camel chymosins were conducted using HIS2 biosensors (Sartorius, 18-5114). All dilutions of antigen and miBd were prepared with MES acetate buffer without Tween 20 (MES-nT) [50 mM MES, 50 mM sodium acetate, 150 mM NaCl, pH 5.5]. Prior to the experiment the HIS2 biosensors were incubated for at least 10 min in MES-nT. For the binding assay, we used the miBd at 200 nM in MES-nT for immobilisation on HIS2 biosensors. Firstly, the affinity towards camel chymosin was measured in seven concentrations ranging from 1,000 nM to 100 nM [1000, 800, 600, 400, 300, 200, and 100 nM, respectively]. Next, after an extended dissociation of 1,200 seconds, the affinity towards the camel chymosin variant was measured employing the same sensors with immobilised miBd A1. The concentration of analyte ranged from 200 nM to 20 nM in the same increments as used for bovine chymosin. Experiments were performed at 30 °C, and data were fitted with a 1:1 binding model.

#### C3 - Bovine Chymosin

The affinity measurements of miBd C3 towards bovine chymosin were conducted using HIS2 biosensors (Sartorius, 18-5114). All dilutions of antigen and miBd were prepared with MES acetate buffer without Tween 20 (MES-nT) [50 mM MES, 50 mM sodium acetate, 150 mM NaCl, pH 5.5]. Prior to the experiment the HIS2 biosensors were incubated for at least 10 min in MES-nT. For the binding assay, we used the miBd at 200 nM in MES-nT for immobilisation on HIS2 biosensors and measured the bovine chymosin as analyte in solution in 2-fold dilution series starting at 1,000 nM and ending in 16 nM. Experiments were performed at 30 °C, and data were fitted with a 1:1 binding model.

#### H2 - Bovine Chymosin, Camel Chymosin (wild type and variant)

The affinity measurements of miBd H2 towards bovine and camel chymosins were conducted on an Octet Red96 (ForteBio) instrument using HIS1K biosensors (Sartorius, 18-5120). All dilutions of antigen and miBd were prepared with MES acetate buffer with Tween 20 (MES-wT) [50 mM MES, 50 mM sodium acetate, 150 mM NaCl, 0.02% (m/v) Tween 20, pH 5.5]. Prior to the experiment the HIS2 biosensors were incubated for at least 10 min in MES-wT. For the binding assay, we used the miBd at 100 nM in MES-wT for immobilisation on HIS2 biosensors. Firstly, the affinity towards camel chymosin was measured in seven concentrations ranging from 1,000 nM to 50 nM [1000, 800, 600, 400, 200, 100, and 50 nM, respectively]. Next, after an extended dissociation of 1200 seconds, the affinity towards the camel chymosin variant was measured employing the same sensors with immobilised miBd H2. The concentration of the analyte camel chymosin variant ranged from 80 nM to 1 nM in seven increments [80, 50, 30, 10, 5, 2, 1 nM]. Lastly, we measured the affinity towards bovine, again employing the same biosensors with H2 loaded. For bovine chymosin we defined a concentration range between 120 nM and 10 nM with seven increments [120, 100, 80, 60, 40, 20, 10 nM]. Experiments were performed at 30 °C, and data were fitted with a 1:1 binding model.

### Functional inhibition – Screening and estimation of K_**I**_^**app**^

Assessment of the capability of the binder candidates to inhibit chymosin in milk was investigated on the basis of an optical activity assay in plate format performed essentially as described in Kappeler et al. (2006)^25^ with some modifications.

Binders and chymosin were diluted in 84 mM acetate pH 5.5 supplemented with 0.2% Tween 80. Chymosin was typically diluted to 0.1-0.2 IMCU/mL (final concentration was determined based on activity with no inhibitors added. The estimated concentrations were: Bovine ∼ 0.23 nM; WT & mutant camel ∼ 0.12 nM; Hedgehog ∼ 0.031 nM) and mixed with 10 µL binder solution in a microtiter plate. The assay was initiated by addition of 250 µL 4% reconstituted skim milk powder (Chr. Hansen, batch Z910600031) supplemented with 10 mM CaCl_2_. and regular measurement (every 20 s) at 880 nm at 32 °C for up to 12 hours.

Quantification of the activity was based on a calibration standard curve of bovine chymosin reference (Chr. Hansen A/S) from 0.0027 IMCU/mL to 0.2 IMCU/mL.

K_I_^app^ was determined using the Morrison approximation for determination of K_i_ under tight binding conditions (Eq.1) through the GraphPad Prism 10.2.1 suite.^4,5,25,31^

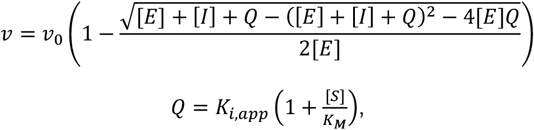

Equation 1 Determination of K_I_^app^ under tight binding conditions

Where v is the reaction rate, v_0_ is the reaction rate without addition of inhibitor, [E] is the concentration of enzyme, [I] s the concentration of inhibitor, and [S] is the concentration of substrate.

A base assumption for the use of this model in the study, is that the relative changes to clotting activity upon inhibitor binding can replace the relative changes to the initial reaction velocity (v).

### Preparing miBd-coated Sepharose matrix for whey filtration

For every coupling reaction (binder), 1 mL NHS-activated Sepharose 4 Fast Flow (Cytiva, 17-0906-01) beads in suspension were transferred to a 5 mL gravity flow column and prepared essentially as described in the instructions for use.

Briefly, the resin was washed 10 times with 1 mL 1 mM HCl to wash away the storage solution. Afterwards, 250 µL of the binder solutions with concentrations from ∼300 µM to 500 µM were mixed with 250 µL coupling buffer (0.2 M bicarbonate, 0,5 M NaCl pH 8,3) and transferred to the drained beads leading to an average concentration on the beads of app. 70 µM to 130 µM assuming full coupling. The beads and binders were incubated for 2 hours on a tube roller at room temperature (∼22 °C). The coupling pH was measured as 7.3.

After binding, excess liquid was drained off and a sample was reserved for SDS-PAGE. The resin was suspended and kept in 0.1 M Tris-HCl, pH 8.5 for 2 hours at room temperature to saturate remaining binding sites. Finally, the resin was washed for 3 cycles with - 3 x 1 mL 0.1 M Tris-HCl, pH 8.5 followed by 3 x 1 mL 0.1 M acetate, 0.5 M NaCl pH 4,5 and considered ready to equilibrate in binding buffer for use.

### Filtration of chymosin in whey by gravitational flow columns

The binding capacity of the binder coupled beads were tested by transfer of 0.2 mL of a 50% freshly prepared bead suspension to a 1 mL gravity flow column. The beads were equilibrated in filtered whey (0.2µm) from Mozzerella pH 6.4 or 84mM acetate (pH 5.8) and drained. 1 mL whey or acetate buffer containing 80 IMCU/mL bovine chymosin was passed through the beads and the flow through was collected. Afterwards the beads were washed with 1 mL acetate buffer followed by 1 mL eluent (either 1 mL 0.1M citrate pH 2.8 or 1mL 0.1M NaOH pH 11). An estimated mass balance was established by determination of the chymosin activity in each of the fractions.

Resin recycling was tested with the miBd A1 coated-resin and 30 IMCU was chosen as the load, since it approached the capacity of the 0.1 mL resin. The beads were treated as above, with 1 mL additions of whey, followed by a 1mL buffer wash, a 1mL elution (0.1M citrate pH 2.8 or 1mL 0.1M NaOH pH 11), followed by an equilibration in 1mL whey. Activity of the FT was monitored for every cycle to determine change in the binding capacity.

