## Supplementary figures and images for "Clearing the whey: Efficient inhibition and removal of chymosin from whey by de novo designed protein-based inhibitors"

### A1_bov_fit_on_processed_data.jpg

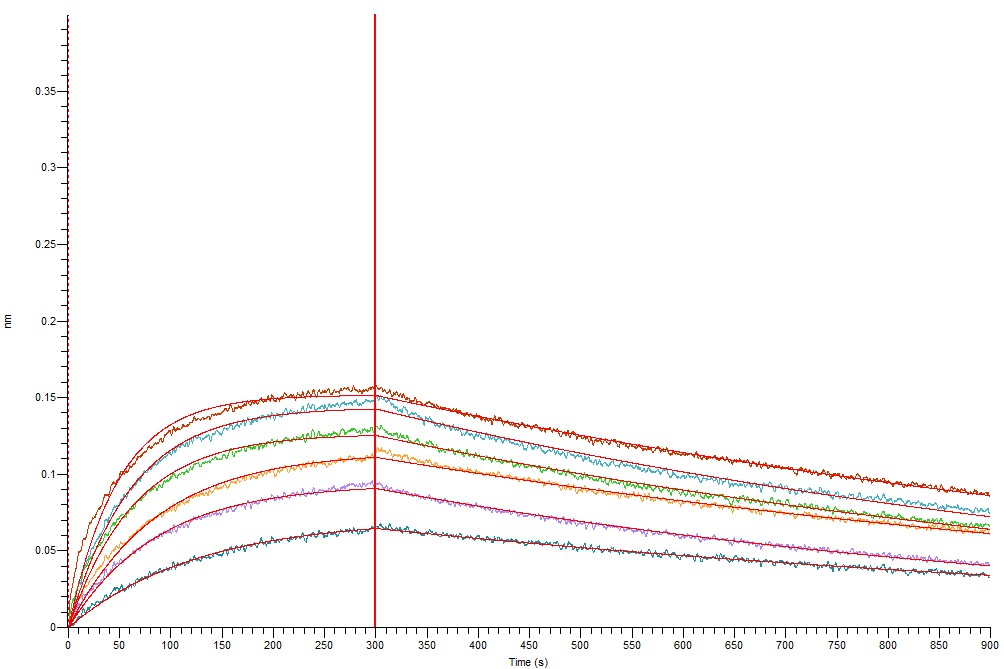

### A1_bov_fit_on_processed_data_2.jpg

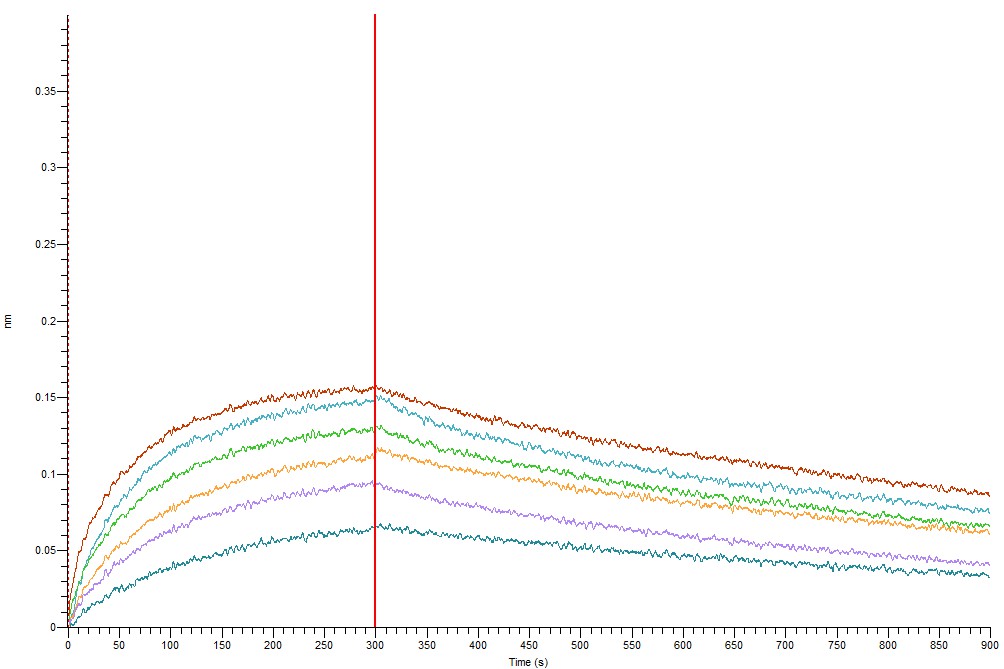

### A1_bov_fit_steady_state_kinetics.jpg

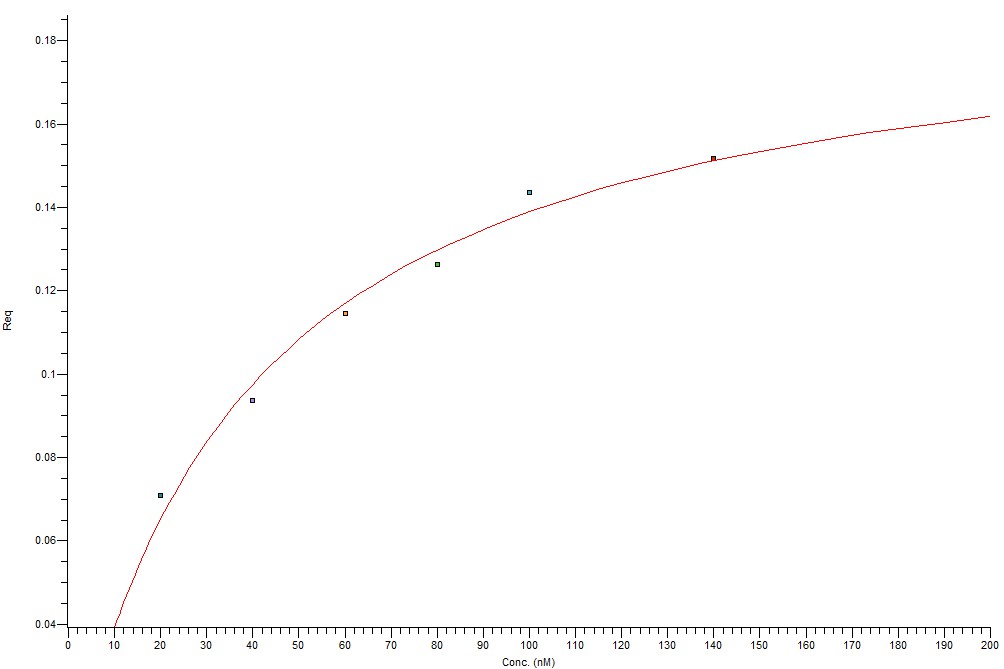

### A1_cam_fit_on_processed_data.jpg

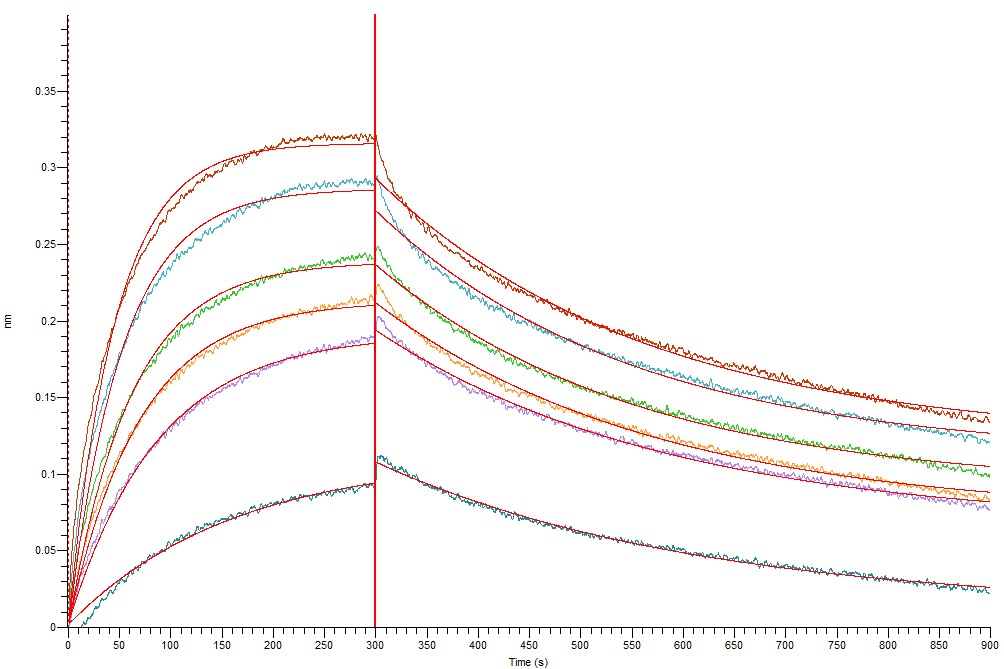

### A1_cam_fit_on_processed_data_2.jpg

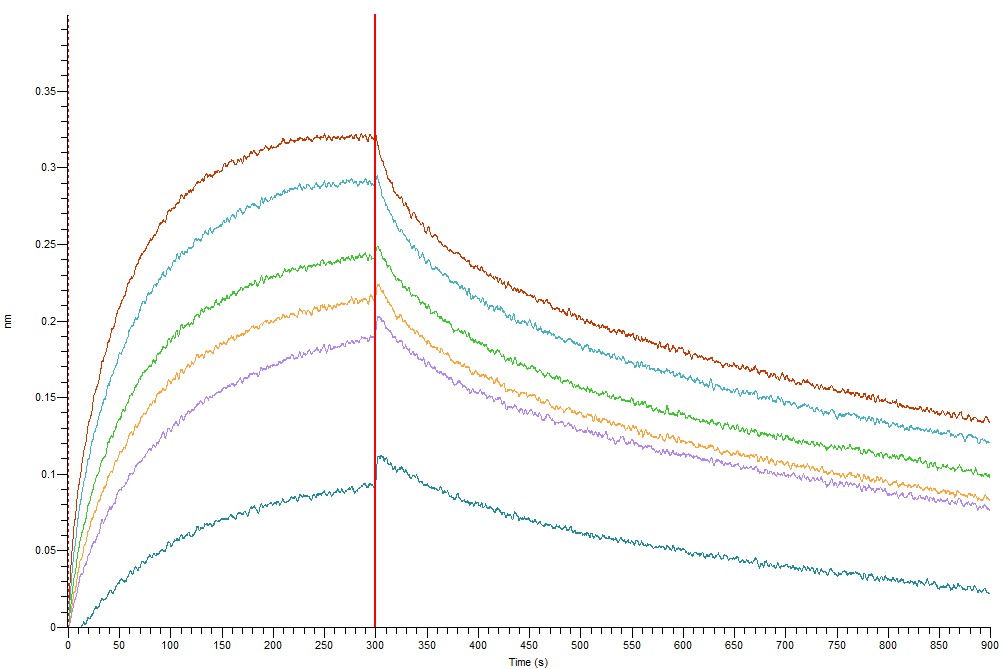

### A1_cam_fit_steady_state_kinetics.jpg

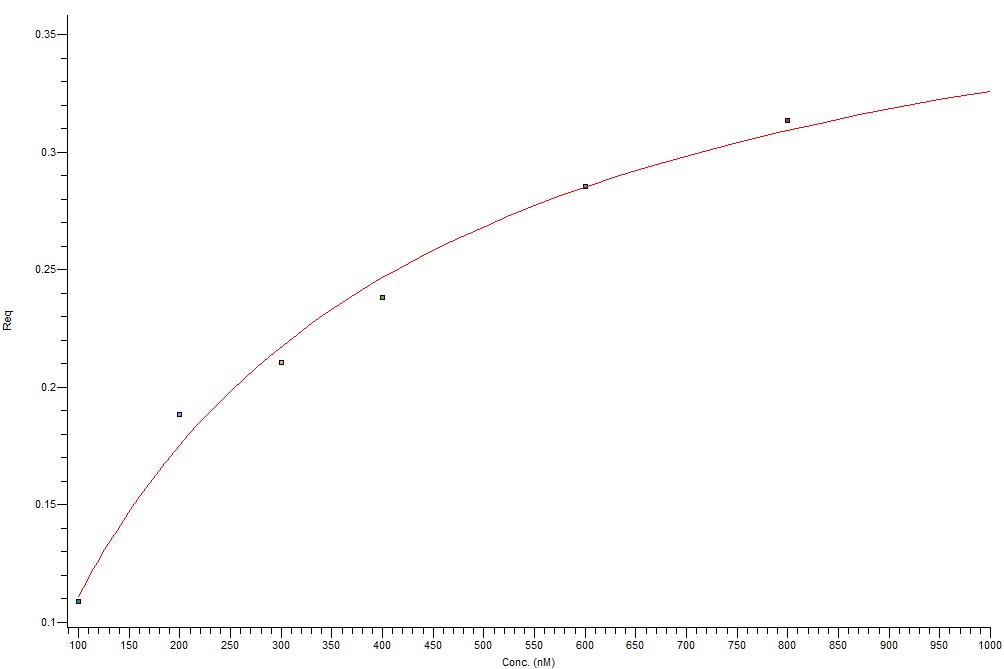

### A1_camS_fit_on_processed_data.jpg

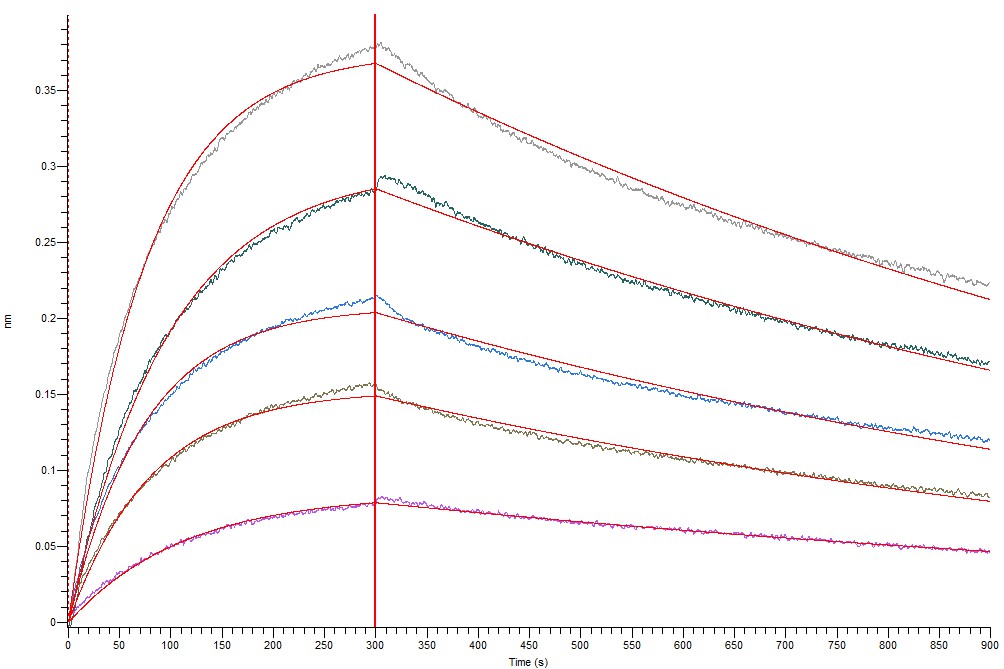

### A1_camS_fit_on_processed_data_2.jpg

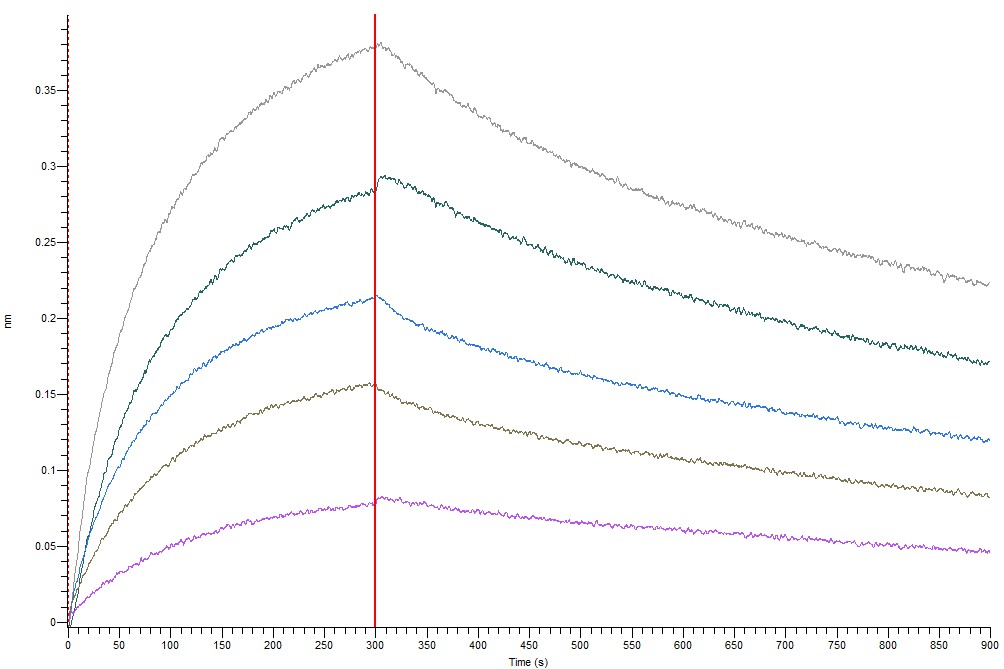

### C3_bov_fit_on_processed_data.jpg

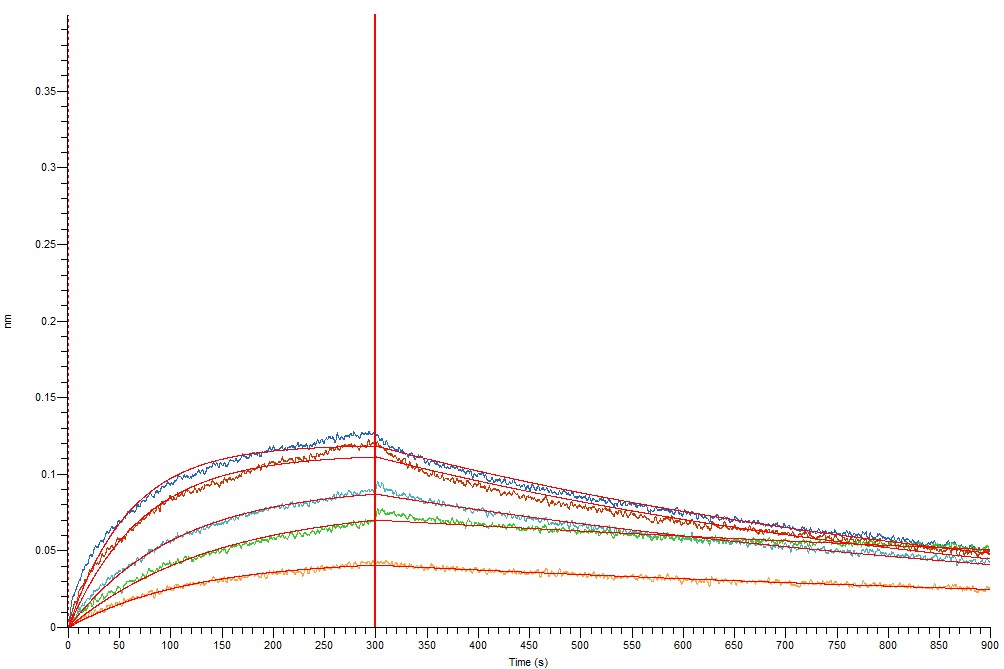

### C3_bov_fit_on_processed_data_2.jpg

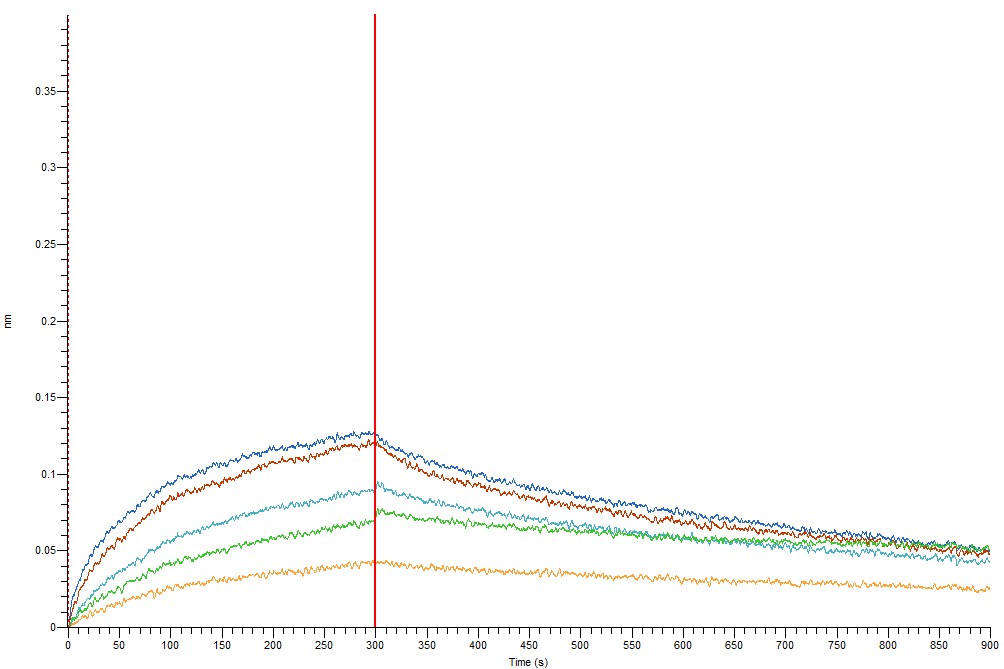

### C3_bov_fit_steady_state_kinetics.jpg

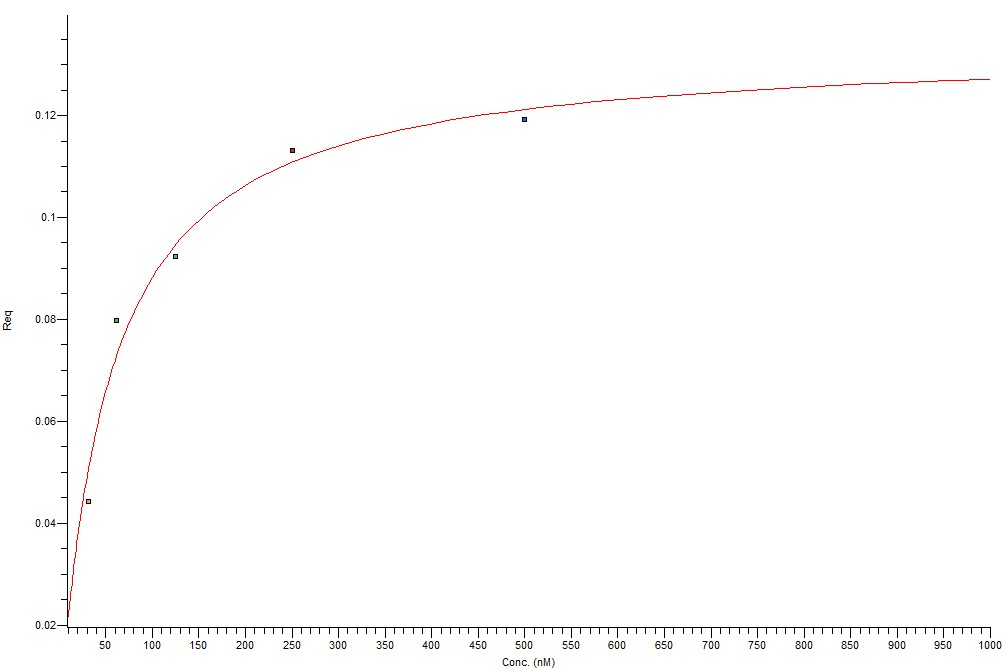

### epiclus_bovine_interaction_plot.png

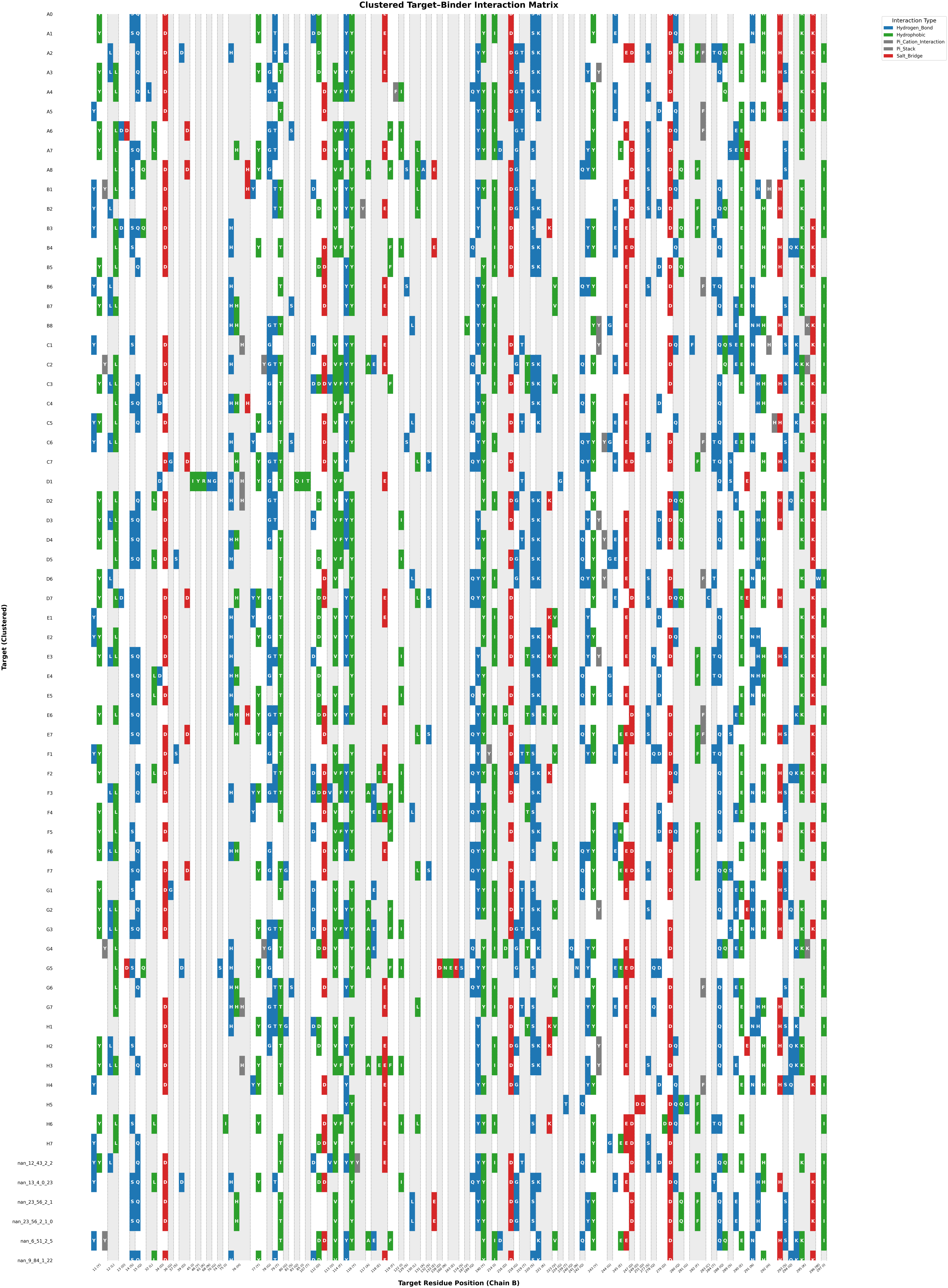

### epiclus_camel_all_miBd_interaction_plot.png

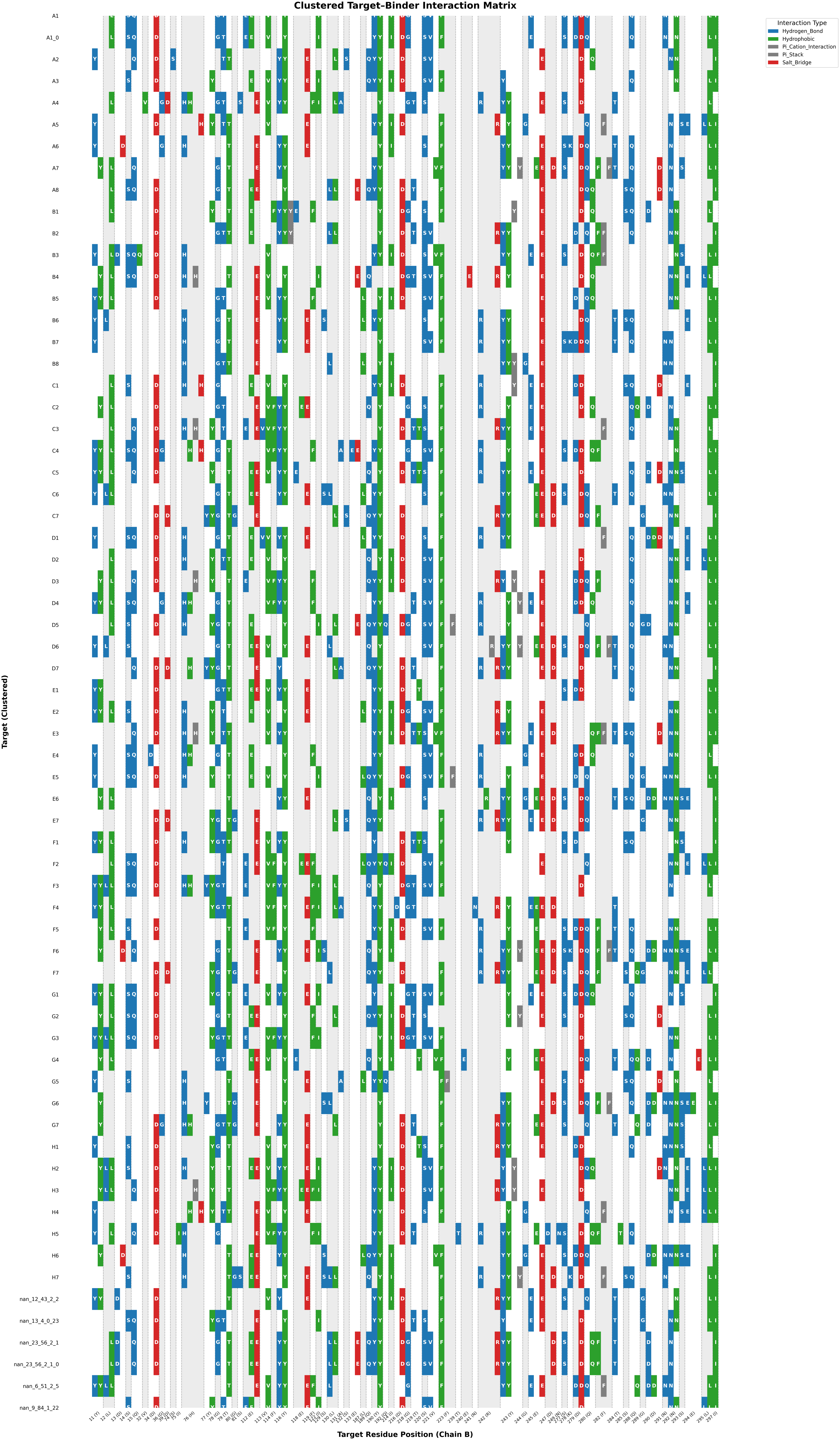

### H2_bov_fit_on_processed_data.jpg

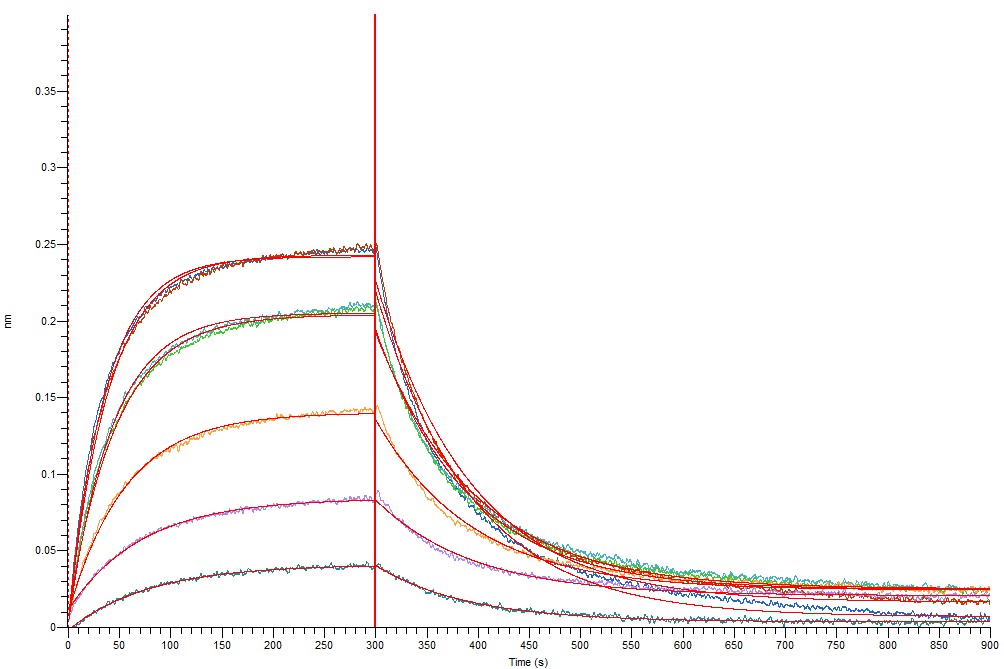

### H2_bov_fit_on_processed_data_2.jpg

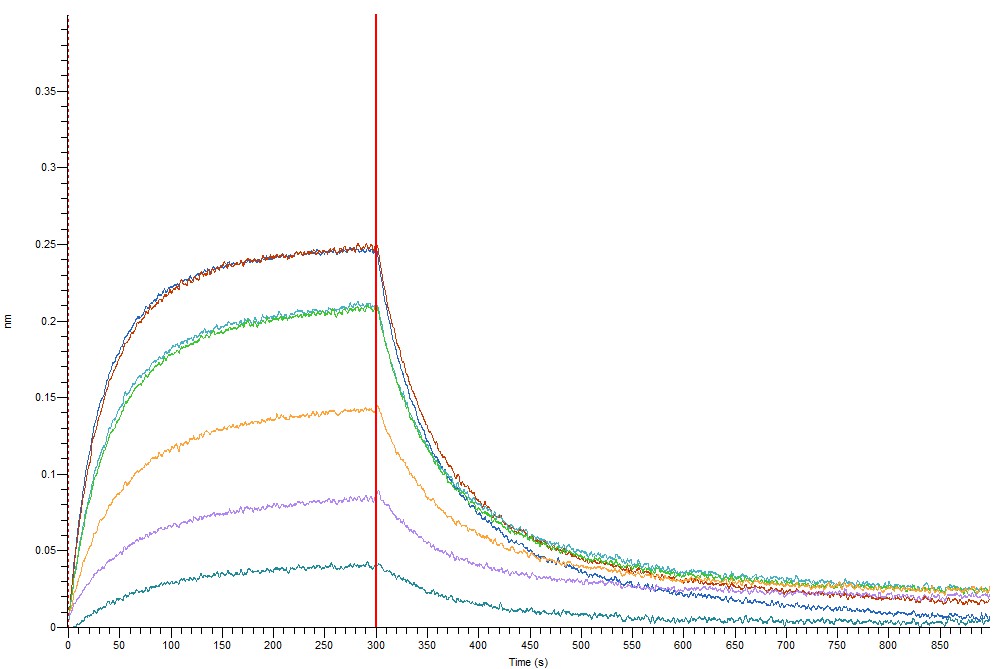

### H2_bov_fit_steady_state_kinetics.jpg

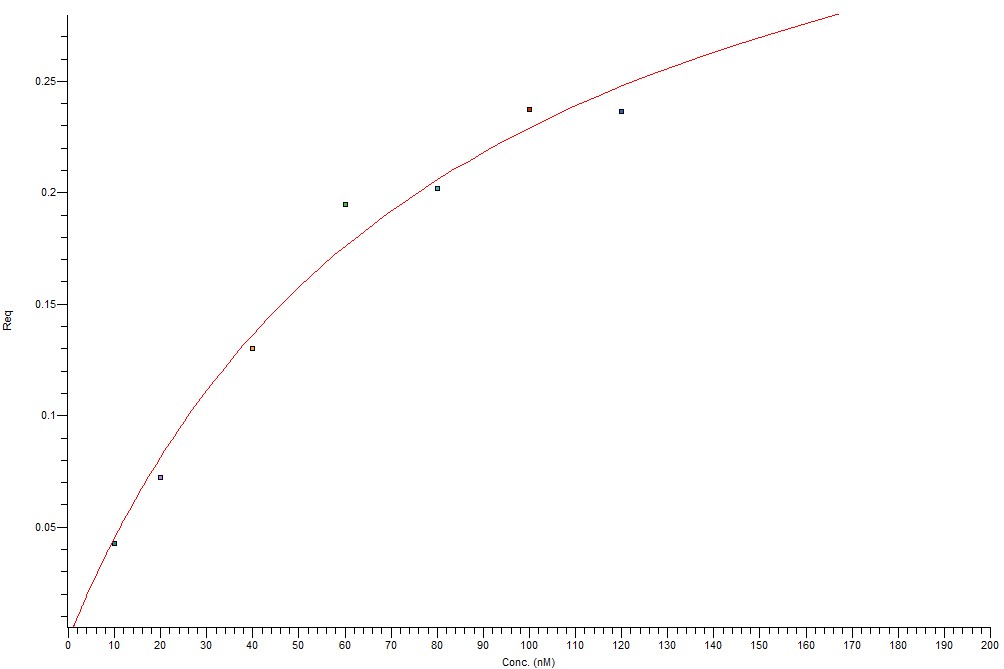

### H2_cam_fit_on_processed_data.jpg

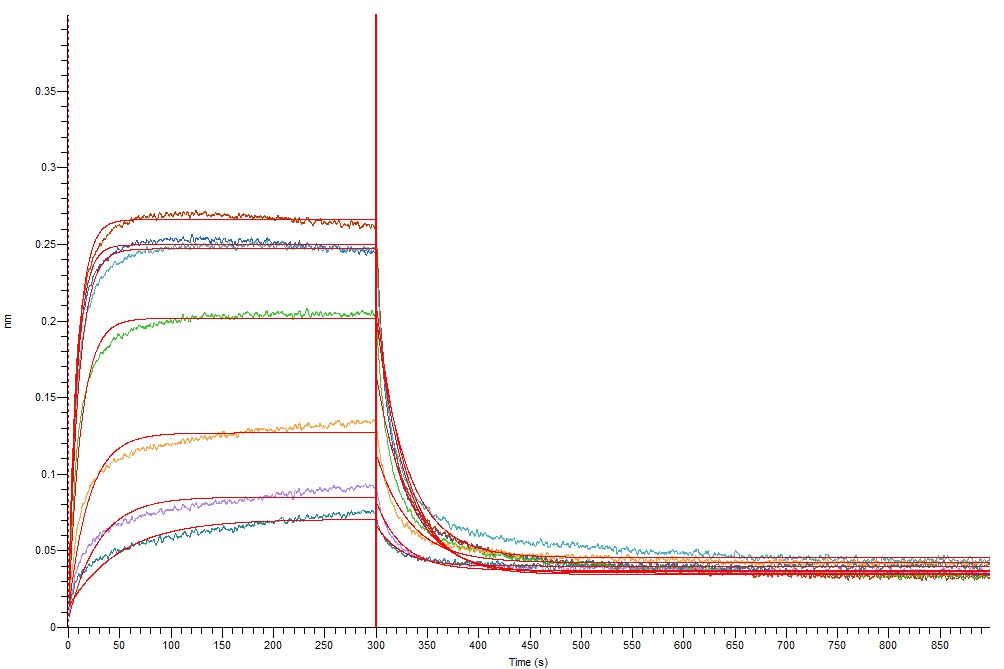

### H2_cam_fit_on_processed_data_2.jpg

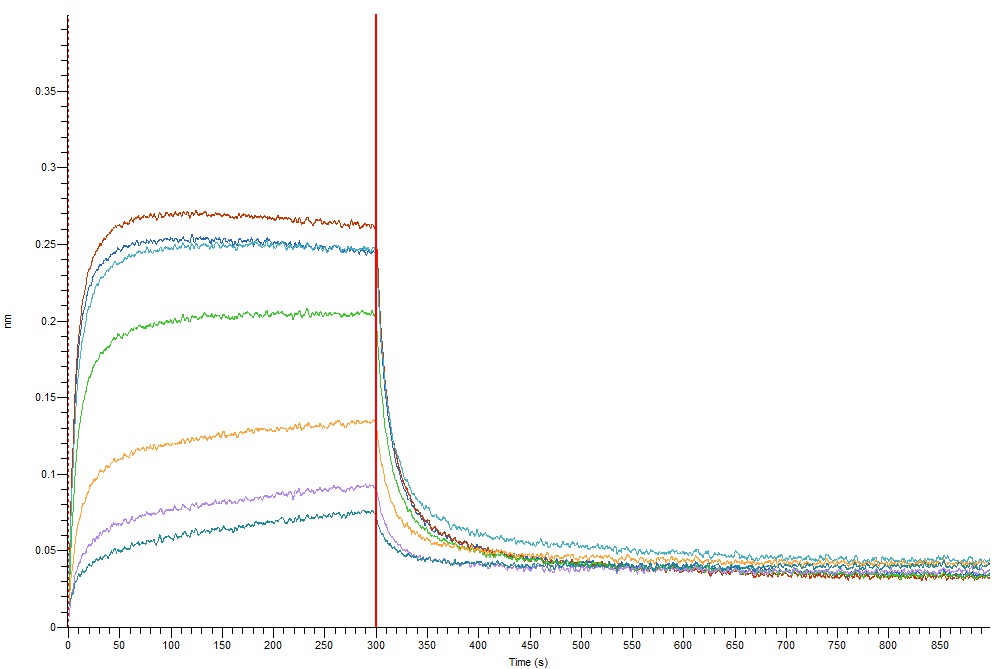

### H2_cam_fit_steady_state_kinetics.jpg

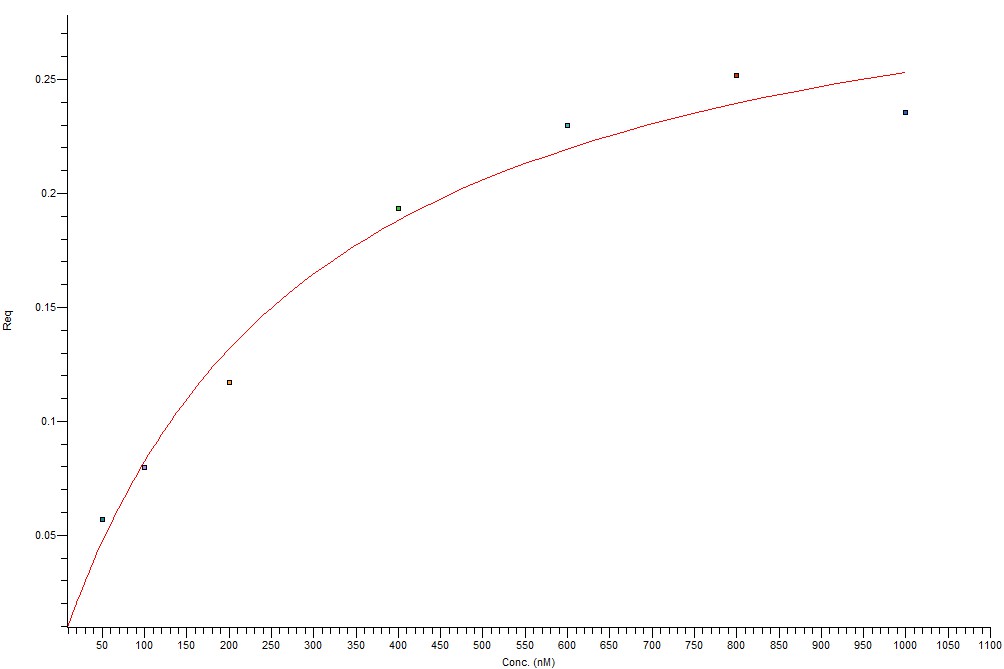

### H2_camS_fit_on_processed_data.jpg

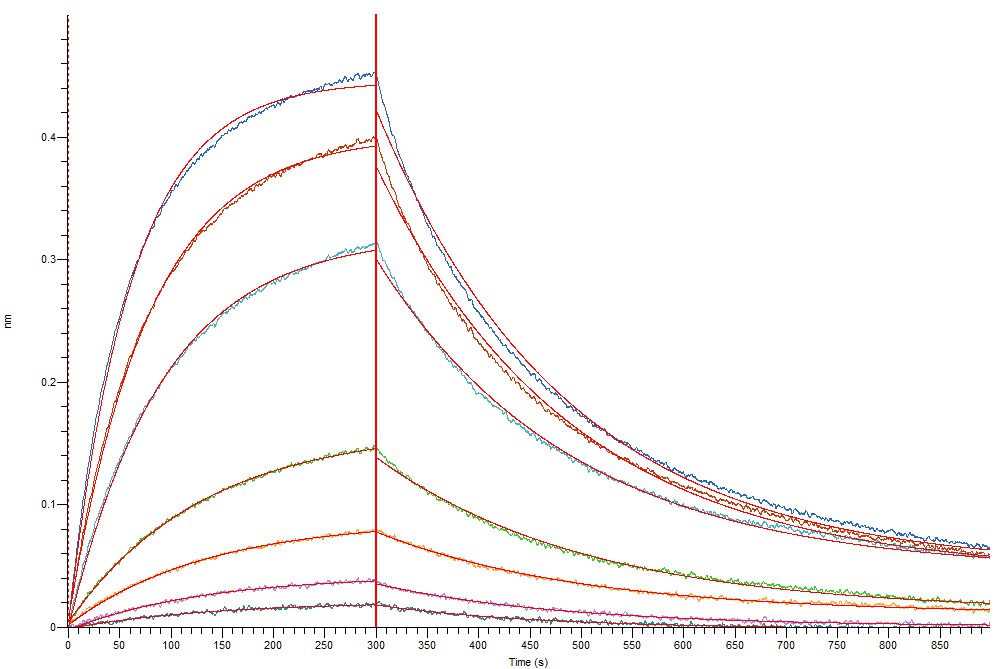

### H2_camS_fit_on_processed_data_2.jpg

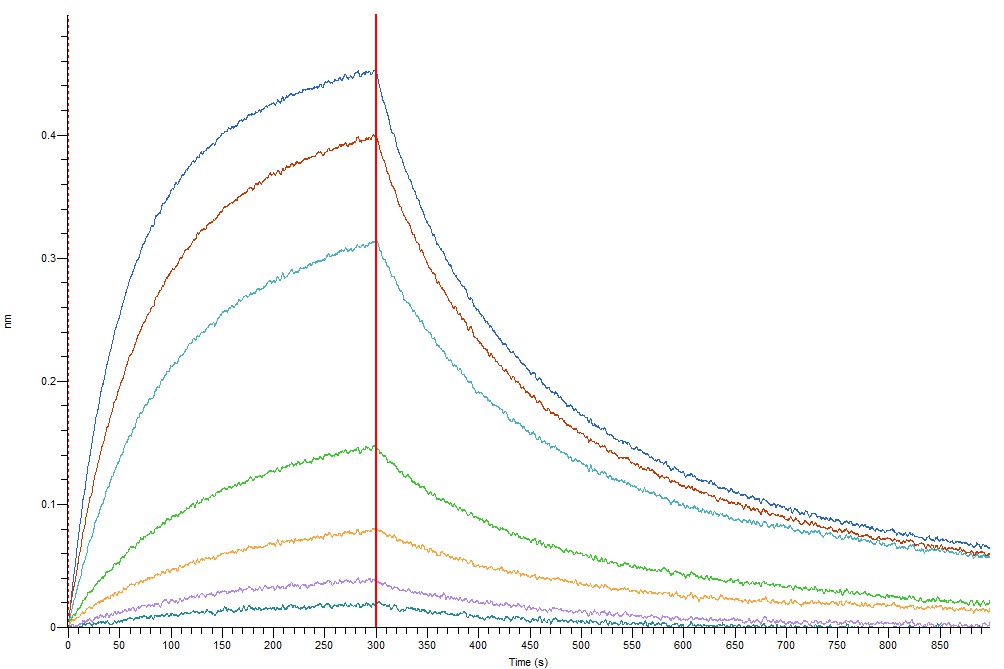

### H2_camS_fit_steady_state_kinetics.jpg

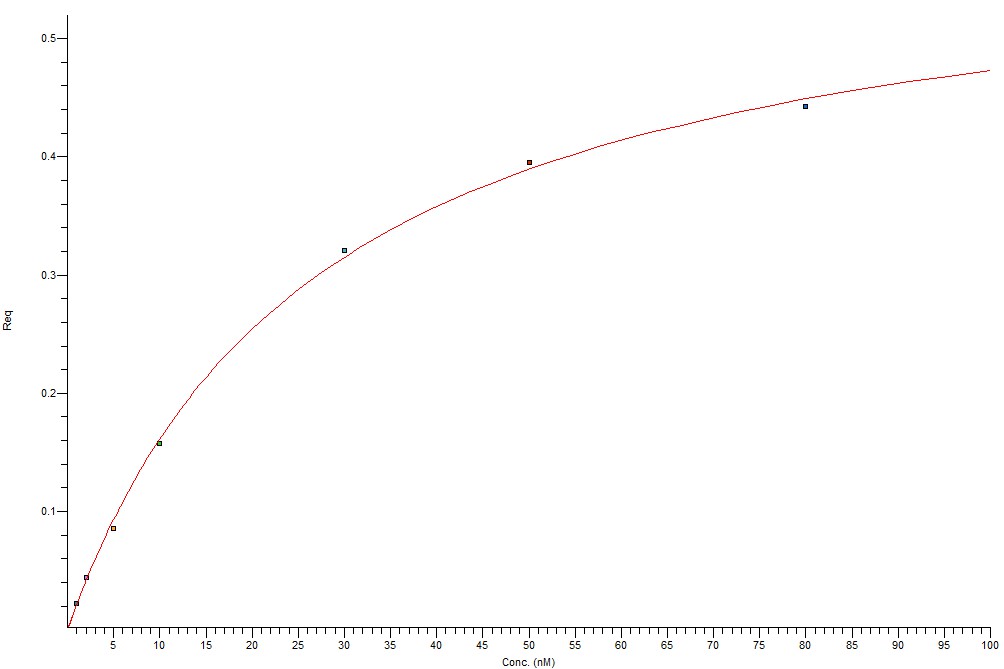
